# Integrating Microchannels and Flows into 3D Printable Granular Hydrogel Matrices

**DOI:** 10.1101/2025.06.08.658465

**Authors:** Emily Ferrarese, Emily Swanekamp, Thuy-vi Bui, Christopher B. Highley

## Abstract

Microfluidic systems incorporating or contained within hydrogels are important in creating microphysiological systems (MPSs). Often naturally derived hydrogels are used, as their inherent bioactivity supports dynamic cellular behaviors. Hydrogel biomaterials that are partly or fully synthetic are desirable in engineering systems with specific, designed properties, though they typically lack bioactive features of natural materials without additional molecular design. In particular, permissive biomaterials enable physiologically relevant dynamic cellular behaviors. Granular hydrogels offer inherent permissiveness, owning to porosity between particles and dynamic behaviors in the absence of interparticle crosslinking. However, applying these in MPS to model tissues requires stable channels to perfuse fluid in these dynamic systems. Here, we establish channels within granular hydrogels to enable perfusion through spatially controlled interparticle crosslinking. Selective crosslinking allowed for the formation of stable channels while allowing the microparticles of a granular hydrogel between two channels to remained uncrosslinked. This allowed spatiotemporal control of signals within an environment established from microparticles without interparticle crosslinking. Fluorescently tagged molecules allowed for the visualization of controlled soluble gradients between two channels within the device. Additionally, embedded 3D printing processes can be used to specify material composition within the system, demonstrating integrated technology for engineering well-defined hydrogel systems. Integrated microfluidic-based control over soluble signals in a system that is compatible with 3D printing processes will establish a basis for building MPSs for broad applications, and the ability to maintain granular systems in culture without interparticle crosslinking will enable design of synthetic hydrogels that access unique dynamic properties within these systems.

## Introduction

Microfluidic technologies have enabled the creation of physiological systems that reproduce complex biology and patient-specific diseases *in vitro* within lab-on-a-chip devices.^1–3^ These microphysiological systems (MPSs) can combine control over flow and control over biophysical and biochemical environments cells experience to model tissues for applications in drug discovery,^4–6^ studying disease progression,^7,8^ and personalized medicine^5,9,10^. As cell decisions are dependent on cells’ integration of soluble and matrix-tethered biochemical signals as well as mechanical environmental cues, designing synthetic systems in well-defined hydrogels in which microflows can deliver soluble cues will present opportunities to understand and perturb physiological systems.^11,12^ Microfluidic devices make it possible to establish gradients or to precisely deliver signals to cells in 2D and 3D environments.^13–15^ Additionally, in establishing vascularized microphysiological models, hydrogels integrated within microfluidic devices can contain endothelial cell linings of microfluidic channels and help drive microvascular maturation.^16–20^

In microfluidic systems, natural hydrogel materials like fibrin or collagen, have been shown to readily support cellular self-organization in multicellular structures, such as microvascular networks formed by vasculogenesis or angiogenesis.^12,21,22^ However, fully- or partially synthetic systems are often desired to achieve designed control over material properties,^11,23^ including through processing by photopatterning and photolithographic 3D printing, which allows for additional control of material compositions in microfluidic devices.^24,25^ In these systems, mesoscale channels can readily be established directly in microfluidic devices by casting around needles^17,26,27^ or molds^28–30^, but microscale vasculature typically requires hydrogel design that permits cells to self-assemble microvascular structures. Vasculogenesis and angiogenesis can be achieved, for example, through crosslinking engineered to degrade^31^, or to be dynamic^32^ in response to cellular activity.

Recently, efforts using granular hydrogel biomaterials have highlighted their strengths in designing MPSs. Granular hydrogels support dynamic cellular behaviour,^33,34^ microvascular structures,^35,36^ and tissue growth^37,38^. Granular hydrogels are formed by packing hydrogel microparticles, whose individual diameters are typically on the order of 10 or 100s of microns, into a bulk scaffold, which results in material systems with unique properties.^39,40^ If microparticle based bulk scaffolds do not include interparticle crosslinking, or annealing,^37^ they can yield and flow in response to mechanical perturbations.^36^ Unlike bulk hydrogels, microparticle-based granular materials are inherently permissive; they support dynamic cellular behaviors such as proliferation, migration, or multicellular self-organization. In granular hydrogels, this is largely a function of micron size pores between individual microparticles. When particles are not crosslinked at their surfaces to one another, they also have the potential to yield and move in response to cellular activities. Granular hydrogels, made from synthetic or semi-synthetic hydrogels, are highly tunable, with control over polymer material, size, stiffness, and degradability for example, that in turn influences cellular behaviors.^35,41–44^ An additional strength of granular hydrogels in designing complex systems is their compatible with bioprinting processes that can be used to specify cellular and material architectures within these materials.^45,46^

Integrating granular hydrogels into devices in which controlled flows could be applied would allow for additional control over microenvironments presented by these materials. In particular, devices that support unannealed granular hydrogels, which lack crosslinking between microparticles, are unstable in standard *in vitro* culture.^39^ Microdevices that can support unannealed granular systems would allow study of cell responses to shear-thinning and self-healing behaviors that are lost with interparticle crosslinking. Additionally, devices might be designed to be compatible bioprinting processes that can be used to specify material complexity in unannealed granular hydrogels. Current examples of devices used to contain granular hydrogels rely on physical features, such as posts to contain uncrosslinked materials.^41,47,48^ As an alternative, perfusable channels embedded within granular hydrogels – which have been achieved in continuous (non-granular) hydrogels by casting the hydrogel around a removeable object^26,49,50^ or material,^27,51,52^ and digital light 3D printing^24^ – have been established within fully annealed granular hydrogels^53–56^, but are unexplored in systems in which unannealed microparticles are preserved within the granular hydrogel volume.

Our approach aims to embed channels fully within the biomaterial, allowing cylindrical channel topographies and avoiding the use of high aspect ratio posts or physical constraints to separate unannealed particles from flow. Here, we designed channels to align with fluidic inlets and outlets, and we use selective crosslinking of granular hydrogels immediately around a channel to stabilize the channel walls in the presence of fluid flow. These allows hydrogel particles that comprise channel walls to be stable in the presence of shear, which might cause unannealed systems to become unpacked and dissociate into the fluid flow. However, in the space between channels, microparticles can remain unannealed (**Figure 1**), allowing for them to retain their dynamic qualities in the presence of perfusive flow. The unannealed granular material can also support 3D printing processes that deposit new materials to create complex architectures within the system. By using two channels to direct flow through the device, it is thus possible to form soluble gradients within the granular hydrogel. Taken together, the system allows for unique control over both soluble cues and unique material-associated cues, the former through control over perfusion and the latter through selective annealing and 3D printing-based control over the placement of material within the microfluidic device.

**Fig 1.**
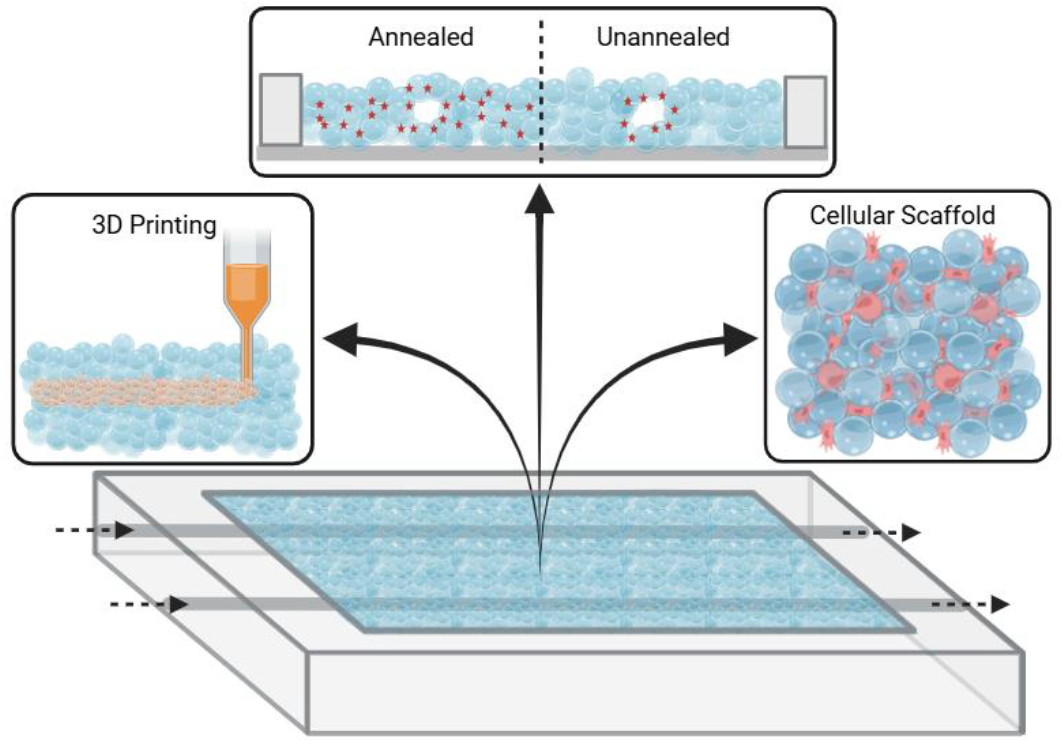
Schematic illustration of a granular hydrogel containing microfluidic device system that could have multiple channels, crosslinked or annealed, between microparticles within the granular hydrogel (indicated by red points in top insert) could be designed throughout the entire scaffold or be localized immediately adjacent to the channels in the hydrogel. With in the volume of the hydrogel between the channels, it would be possible to further define or modify the material structure via 3D printing. Cells might be included among the hydrogel microparticles, if desired.

## Materials and methods

### Norbornene modified hyaluronic acid (NorHA) synthesis

NorHA was synthesized as previously described.^57^ Briefly, HA-TBA was formed by dissolving sodium hyaluronate (Lifecore, 60 kDa) in DI water and mixed with Dowex 50X proton exchange resin for 2 hours. The resulting solution was titrated with tetrabutylammonium hydroxide to pH 7, frozen and lyophilized. HA-TBA was then modified via amidation with 5-norbornene-2-methylamine, anhydrous dimethyl sulfoxide (DMSO), and benzotriazole-1-yl-oxy-tris-(dimethylamino)-phosphon-ium hexafluorophosphate (BOP) under nitrogen at room temperature for 2 h. The reaction was quenched with cold water, purified via dialysis (SpectraPor, 6–8 kDa MWCO) for 3 days with DI water and NaCl, then another 4 days with DI water, frozen and lyophilized. The degree of modification was 15.5% determined by 1H NMR (Supplemental Figure S1).

### Hydrogel fabrication

A NorHA hydrogel precursor solution was made by combining NorHA (3wt%, 0.6 thiol:norbornene (SH:Nb)), with dithiothreitol (DTT, ThermoFisher) photoinitiator lithium phenyl-2,4,6-trimethylbenzoylphosphinate (LAP, 25 mM, Sigma-Aldrich), and deionized water. NorHA hydrogel microparticles were formed using a batch emulsification technique similar to previously reported approaches (**Figure 2**).^43^ Briefly, the aqueous solution was added to light mineral oil with Span 80 (0.5 vol%, Sigma-Aldrich) at a 20:1 oil to aqueous ratio of the oil to aqueous volumes and mixed on a stir plate at 600 rpm. After 1 minute the microparticles were crosslinked under UV light (320-390 nm) at 60 mW/cm^2^ for 5 minutes. The particles were centrifuged at 3,500 rcf for 2 minutes to remove the majority of the oil and surfactant. The particles were then washed sequentially with 2 wt% Pluronic F-127 (Sigma-Aldrich), and 70 vol% ethanol two times. Microparticles were stored in phosphate buffered saline (PBS) at 4°C until use.

**Fig 2.**
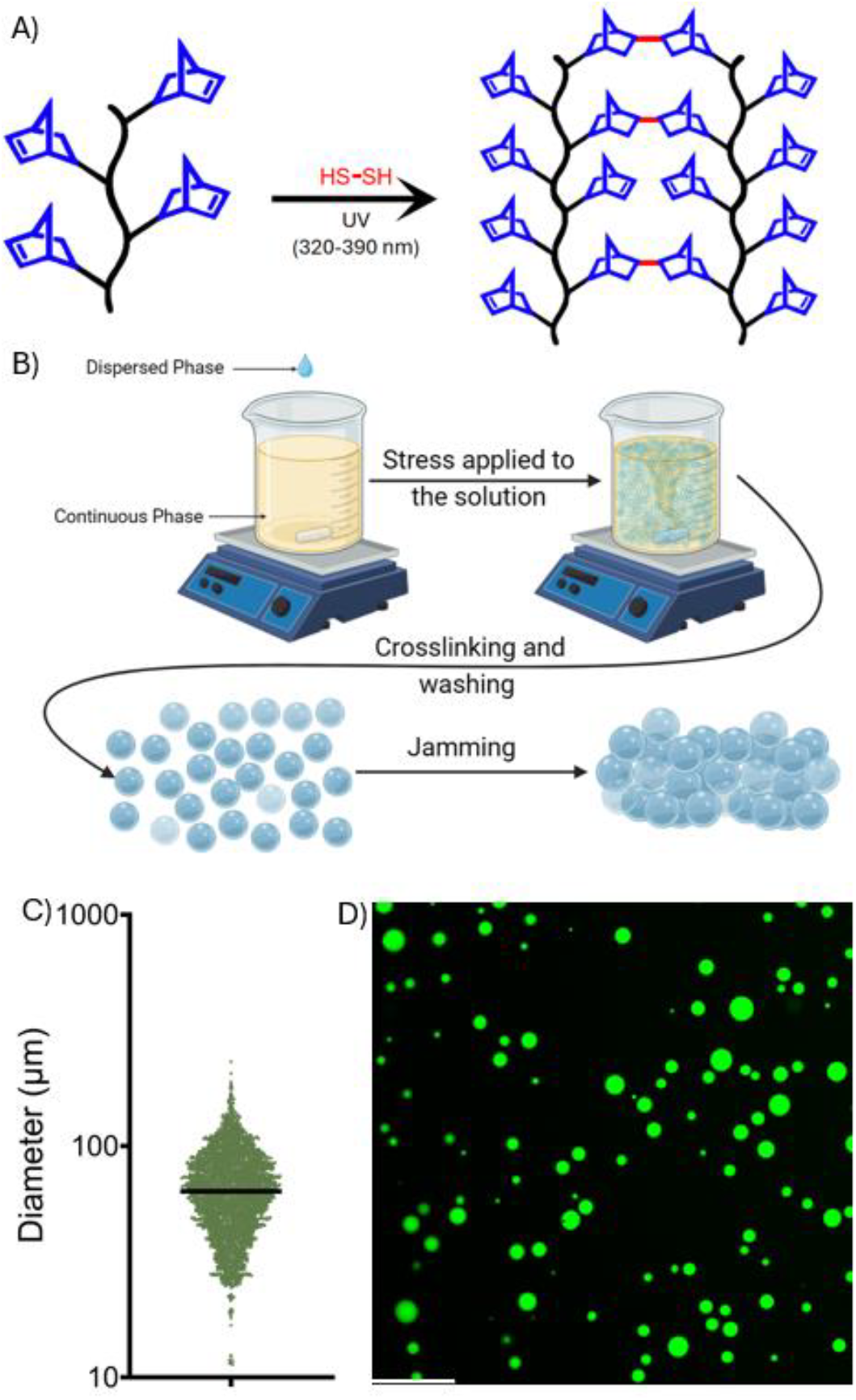
A) Hyaluronic acid (black) functionalized with norbornene groups (blue) and reacted with dithiol crosslinker (red) to form the hydrogel polymer network. B) Batch emulsification technique to fabricate hydrogel microparticles. C) Size distribution and D) representative image of NorHA hydrogel microparticles (scale bar 500µm).

Polyethylene glycol (PEG) microparticles were generated via aqueous two-phase suspension, based off previous approaches.^58^ Briefly, eight arm PEG-norbornene (3wt%, PEG-Nb, 20kDa, JenKem), four-arm PEG-thiol (0.6 SH:Nb, 10kDa, JenKem), and LAP (10mM) were combined in aqueous solution. A 25wt% solution of dextran (70 kDa, Sigma-Aldrich) was mixed with PEG-Nb hydrogel precursor solutions at a 4:1 ratio of continuous (dextran) to dispersed (PEG-Nb) phases. This mixture was mixed at 200rpm for 1 minute prior to crosslinking under UV light at 30 mW/cm^2^ for 5 minutes. Following crosslinking, the resulting particles were suspended in 15x volume of PBS to thermodynamically favor a single-phase solution and centrifuged twice to remove dextran and other unreacted materials.

After microparticles were washed and prior to forming a bulk granular scaffold, all microparticles were allowed to swell overnight in PBS. To form granular hydrogels, microparticles were then packed together by centrifuging a suspension of microparticles at 21,000 rcf for 5 minutes and excess PBS was removed, creating packed microparticles. To allow for photoinitiated interparticle crosslinking, microparticles were resuspended in a 10wt% PEG-SH and LAP (25mM) crosslinking solution prior to centrifuging. Packed microparticles were loaded into an 8 mm diameter biopsy punched polydimethylsiloxane (PDMS) slab that was 5 mm think and placed under UV light for 5 minutes at 100 mW/cm^2^ to crosslink microparticles together. Granular hydrogel disks were placed in PBS at 37°C for 24 hours to quantify stability.

### Hydrogel characteristics

Fluorescein isothiocyanate-dextran (FITC-dextran, 1mg/mL, 2MDa, Sigma-Aldrich) was added to the hydrogel precursor solution to visualize the microparticles using fluorescent microscopy in order to characterize microparticle size. Microparticles were suspended in PBS and imaged using Lecia DMi8 fluorescent widefield microscope.

Rheological behaviors of packed microparticles were characterized using oscillatory shear rheology (DHR3 rheometer, TA instruments). Pack NorHA microparticles were analyzed using a 20 mm parallel plate geometry set at a gap ten times the average particle diameter. A solvent trap was used to reduce evaporation. Cyclic stain sweeps alternating between high (500%) and low (1%) strain, were used to assess the yielding and recovery responses of the packed microparticles.

### Device fabrication

A computer-aided design (CAD) model of the negative mold of the device was made using Autodesk Fusion 360 software and printed using stereolithographic 3D printer (Form 2, Formlabs). The mold was washed with isopropanol twice then cured for 30 minutes at 75°C. 27-gauge needles were inserted through the negative mold in designed locations, and PDMS (Sylgard 184, Dow Corning) was added into the reverse mold. PDMS was cured at 37°C for 24 hours. The PDMS device frame was then removed from the mold, plasma treated with air, and bonded to a glass coverslip (#1, 60×24mm, VWR).

### Device loading

27-gauge needles were placed through the channels of the PDMS device to template for channels within the hydrogel. To form the continuous NorHA hydrogels within the device, NorHA precursor solution was added and crosslinked under UV light. Alternatively, to form a fully annealed granular hydrogel within the device, microparticles were resuspended in the crosslinking solution, then centrifuged at 21,000 rcf for 5 minutes. These packed microparticles were loaded into the PDMS device with 27-gauge needles in place and crosslinked under UV light.

To form unannealed granular hydrogels containing channels, interparticle crosslinking was desired only immediately adjacent to the channels, leaving the bulk of the microparticles uncrosslinked to one another. To achieve this, LAP was localized to the needles used for templating the channels, and microparticles were resuspended in a solution containing 10 wt% PEG-SH without LAP, packed via centrifugation, and then added to the device with 27-gauge needles in place. LAP was localized to the needles by pipetting a 25 mM LAP solution over the needles after the needles were positioned within the PDMS frame and before adding the packed microparticles. Before adding packed microparticles to the device, the LAP solution was aspirated, leaving needles surfaces wetted with LAP solution. Upon adding the packed microparticles to the device, crosslinking was initiated under UV light.

In conjunction with all hydrogels used – the continuous hydrogel, the fully-annealed granular hydrogel, and the unannealed granular hydrogel – a glass coverslip (#1, 18×18 mm, VWR) was placed on top of the device immediately after loading to prevent hydrogels from drying out. Then the device was flipped and placed under UV light at 100 mW/cm^2^ for 5 minutes to crosslink the hydrogel. Then needles were slowly removed to ensure the channels were not disrupted.

### Diffusion characteristics

Devices containing a hydrogel with channels were perfused with fluorescently labeled albumin (FITC-albumin, 50 µg/mL, Sigma-Aldrich)^59^ through a channel at a flow rate of 1 mL/hr using a syringe pump for 25 minutes. Timelapse fluorescent and brightfield images were taken using Leica DMi8 widefield microscopy at 0, 5, 10, 15, 20, and 25 minutes.

For dual-channel diffusion experiments, a rhodamine B (RhodB, 12 µg/mL, Sigma-Aldrich) solution was prepared. The device was prepared with PEG-Nb packed microparticles and selectively annealed along the channel. Once the needles were removed FITC-albumin and RhodB were introduced in solutions flowing through different parallel channels in opposite directions. Timelapse images were taken every 5 minutes of the whole device. Images including RhodB were pseudo-colored blue in figures for accessibility.

### Fluorescent micro particles

Devices loaded with unnannealed NorHA microparticles were perfused with medium containing suspensions of fluorescent microspheres (FMs, FlouSpheres Carboxylate-Modified Microspheres, 10% vol/vol, ThermoFisher). FMs were 1µm in diameter and perfused at a flow rate of 1 mL/hr. Timelapse images were taken over 5 minutes.

### 3D Printing

A biomaterial ink for 3D printing was formed from a granular hydrogel^60^, here based on gelatin microparticles. Gelatin microparticles were formed via a batch emulsification technique as previously described.^61^ Briefly, a 15 wt% solution of gelatin type B (Sigma-Aldrich) with India ink (1:1000) was made and warmed to 80°C to dissolve the gelatin. The warmed gelatin solution was added to a warmed (80°C) light mineral oil solution containing 2 vol% Span 80 and emulsified via homogenized at 2,000 rpm for 3 minutes, before being placed in a 4°C refrigerator to cool. The resulting microparticles were washed sequentially with 2 wt% Pluronic F-127, and PBS 5 times. Gelatin microparticles were packed together at 21,000 rcf for 5 minutes to create a granular gelatin hydrogel to be used as an ink in 3D printing. A 3D printer equipped for volumetric extrusion (BIOprinter, FELIXprinter) was used print the granular gelatin hydrogel. The granular hydrogel was loaded into a syringe that was placed in the printhead for extrusion. Discrete voxels were printed into the center of unannealed NorHA packed microparticles contained within the region in the center of the device between two channels formed, as described above, by templating with a needle. A sample G-code is shown in the supplemental methods.

### Image and Statistical Analysis

Brightfield images of the channel were analyzed to quantify channel diameter using ImageJ. Fluorescent images of diffusion were also processed and analyzed using ImageJ. To quantify diffusion from fluorescent images, a 1 mm distance was traced from the channel boundary perpendicularly into the surrounding hydrogel in the device. Fluorescent intensities along this 1 mm trace were acquired, and background fluorescence was subtracted from each point. Each fluorescent intensity value was then normalized to the maximum fluorescent intensity value of the FITC-albumin solution in the channel. Based on the changes in fluorescence along the trace over time, effective diffusivities within the hydrogel systems were calculated.

Analysis of effective diffusivity included simplifying assumptions.^62,63^ First, it was assumed that the resistance to mass transfer in the liquid is negligible and convective transport axially in the channel ensured a constant concentration of the fluorescent solute in the channel. The concentration at channel wall at the interface with the hydrogel was thus considered constant. Furthermore, over the time course used in experimentation it was assumed that mass transfer was symmetric in all directions, and that the hydrogel was a semi-infinite medium in the radial direction perpendicular to the channel axis.

To approximate diffusivity from observation, the intensity profiles were fitted to Frick’s 2^nd^ law of one-dimensional diffusion:

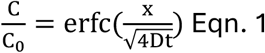

where erfc is the complimentary error function, with FITC-BSA concentration (C), constant FITC-BSA source concentration (*C*_0_), position (x), diffusivity (D), and time (t). The intensity profiles were fitted to Eqn. 1 using curve fitter in MATLAB. Each diffusion experiment performed had n=3 trials. GraphPad Prism 9 was used for all statistical analysis. Data are reported as mean ± standard error of the mean, and particle diameter is reported as mean ± standard deviation. Statistical significance was determined by a one-way ANOVA followed by a post hoc Tukey’s Honest Significant Difference test. Data were graphed with the mean and error bars representing the standard deviation.

## Results

### Hydrogel fabrication and characterization

Hydrogels were crosslinked, both in continuous hydrogels and in hydrogel microparticles as well as at particle-particle surfaces, through a thiol-ene photoinitiated click chemistry reaction (Figure 2A).^64^ Stoichiometric control over NorHA and DTT allowed unreacted norbornene groups to be left after crosslinking for future interparticle bond formation, or annealing. The NorHA microparticles generated by the bulk emulsification process used here were polydisperse with a mean diameter of 66.94±27.32 µm (Figure 2B-C). After centrifugation to form a granular hydrogel, the bulk granular hydrogel system exhibited both viscoelastic properties when the particles were not crosslinked to one another. Under static conditions, it exhibited solid-like properties. Under increasing strains, the bulk material yields, exhibiting liquid-like flow. This strain-dependent behavior in the uncrosslinked system could be observed using oscillatory rheology (Figure 3A). The bulk granular hydrogel formed from packed NorHA microparticles had a storage modulus of 119.1±10.48 Pa and loss modulus of 44.13±8.48 Pa (Supplemental Figure 4), and the difference in the storage modulus compared to a continuous hydrogel (Supplemental Figure S4C) was statistically significant. In the granular hydrogel system, cyclic strain sweeps that alternated between applying low (1%) and high (500%) oscillatory strain demonstrated that the unannealed granular hydrogel had a shear thinning and self-healing behavior (**Figure 3A**) as a bulk material, which is an important property when developing a support material for 3D printing.

**Fig 3.**
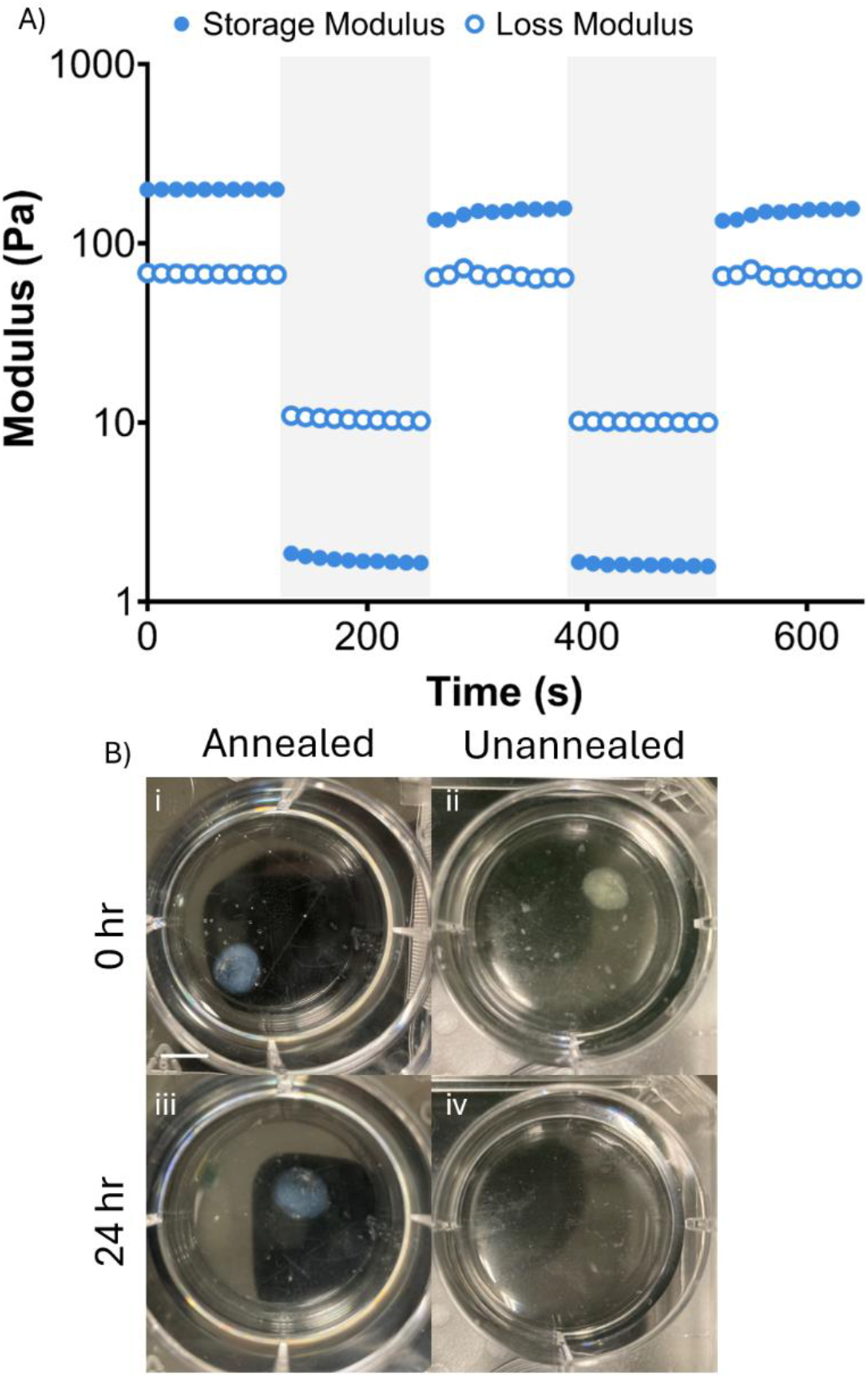
Granular hydrogels display dynamic behaviors in the absence of interparticle crosslinking. A) Cyclic strain sweep alternating between high (white) and low (grey) strain of packed NorHA microparticles, showing the rapid transition from solid to liquid like behavior and back when high strain is applied and removed. C) Packed microparticles that are annealed (i) and not annealed (ii) together right after exposure to UV light and annealed and unannealed microparticles after 24hr (scale bar 5mm).

**Fig 4.**
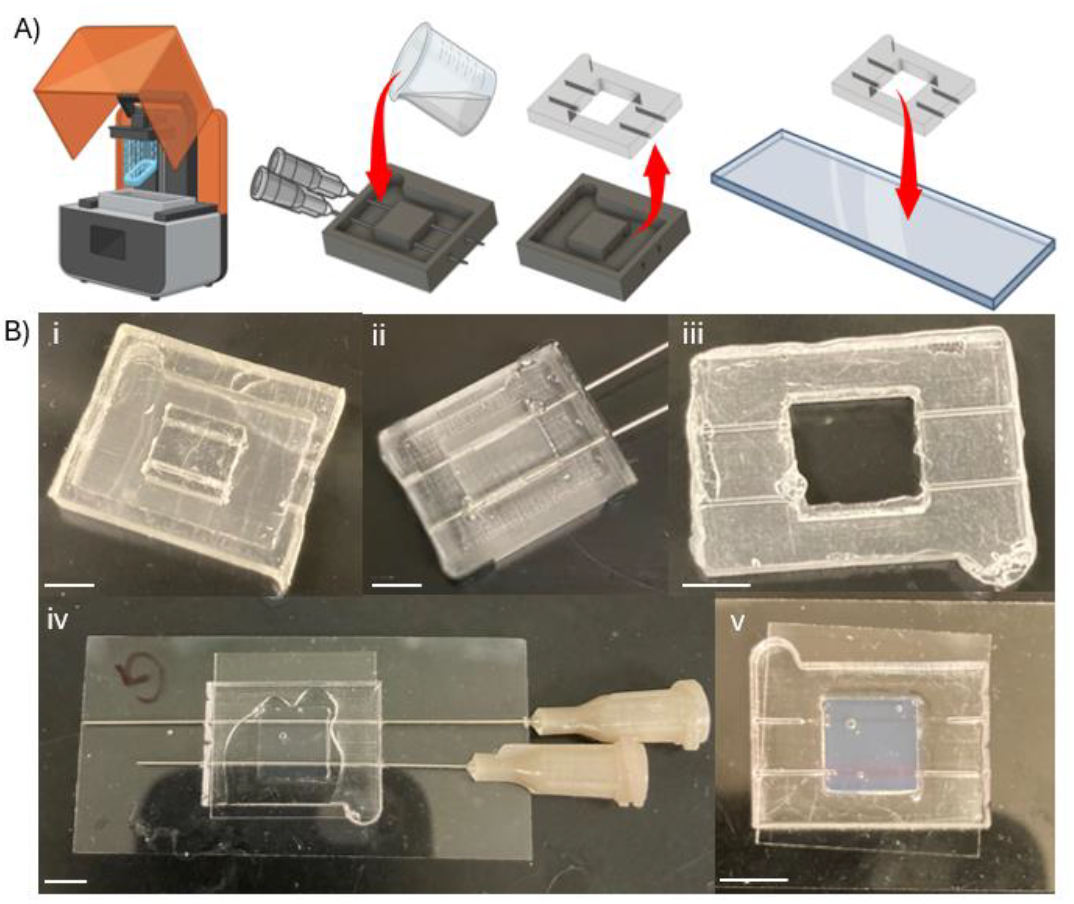
Hydrogels were integrated into PDMS device formed using 3D printed molds. A) The process for manufacturing process of fabricating the microfluidic device was to print a negative mold, insert needles, cast PDMS, and bond PDMS to a glass coverslip. B) Images of key components form the process:(i) the 3D printed negative mold of the device; (ii) the mold loaded with PDMS; (iii) the PDMS device prior to hydrogel loading; (iv) the PDMS device loaded with hydrogel solution prior to; and (v) the device after RhodB injected into channels for visualization (scale bar 5 mm).

When microparticles within a granular hydrogel are annealed they lose their ability to flow in response to an applied stress and to present a dynamic environment, but they are stable against erosion. When granular hydrogels were formed as disks, in absence of a support device, annealed granular gels remained intact after 24 hr whereas granular hydrogels that are unannealed rapidly dissociated as microparticles eroded into the surrounding medium over 24hr (Figure 3B). This highlights the key challenge this work sought to address establishing technology that allows uncrosslinked granular systems to be maintained and perfused over time, towards supporting eventual studies of cellular response to these environments.

### Device Characterization

A 3D printed negative mold was used to fabricate the PDMS device (**Figure 4**) that supports granular hydrogels (Figure 4B, iv) with integrated microchannels (Figure 4B, v). Channels within the PDMS were designed to allow the inlet and outlet fluidic connections to introduce controlled microflows into the hydrogels. As described above, granular hydrogels were introduced into the device and either fully annealed throughout the granular hydrogel bulk or annealed around the channels only, but localizing photoinitiator to the needles. Channel integrity was confirmed via imaging and the introduction of fluidic flows. After removing the 27-gauge needle, which had an outer diameter of 413 µm, brightfield images showed the channels had diameters of 358.3±7.19 µm, 324.7±15.11 µm, and 295±12.94 µm for the bulk, fully annealed, and selectively unannealed hydrogels, respectively (**Figure 5**). A slight statistical significance in channel diameter was observed between only the selectively annealed hydrogel group and the bulk hydrogel.

**Fig 5.**
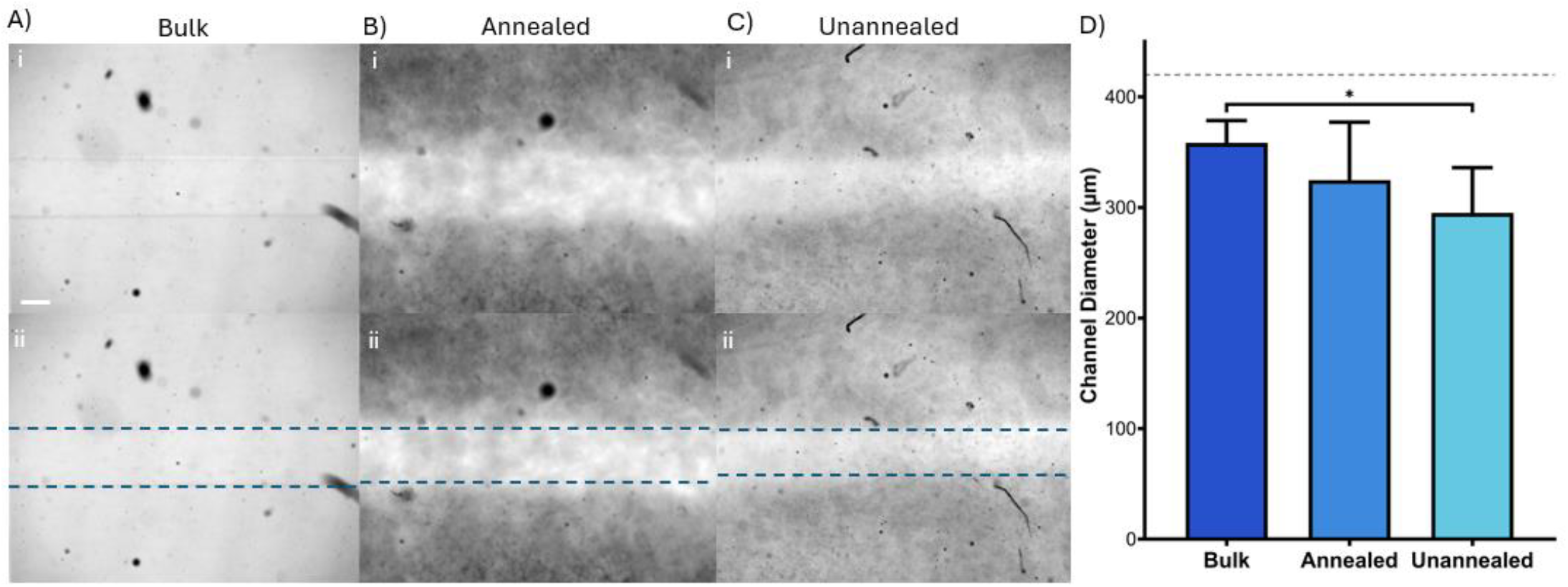
Channels are visualized within hydrogels after crosslinking and removal of the needle using to define hydrogel channel widths. Channels are formed within A) a bulk nanoporous hydrogel, B) a fully annealed granular hydrogel, and C) a granular hydrogel in which crosslinking occurred only among microparticles adjacent to the channel’s edge. The top row shows the brightfield images and the bottom row includes a blue dash line represents the channels boundary. (scale bar 200µm) D) Quantification shows similar channel diameters, with a slight decrease, attributed to increasing in swelling, as crosslinking within the bulk decreases across groups. A statistically significant difference was observed between the channel diameter in the bulk and unannealed hydrogel group. The grey dash line represents the needle diameter. (* p ≤ 0.05)

Medium containing fluorescently tagged albumin (FITC-albumin) that was introduced into the channels allowed visual confirmation that channels remained perfusable during diffusion experiments. FITC-albumin also allowed visualization of protein diffusion from the channel^65^ with time lapse images showing increasing radial distribution of FITC-albumin’s fluorescent signal in the surrounding hydrogel (**Figure 6**). Diffusion from the channels in granular gels – both the fully annealed and unannealed with selective crosslinking - into the surrounding material was compared between annealed and unannealed systems and to diffusion in traditional continuous hydrogels. As expected, the diffusion of FITC-albumin into continuous hydrogels was slower than diffusion in granular systems (Figure 6D), due to the presence of microporosity in the granular systems. The continuous hydrogel consists only of a nanoporous polymeric network outside of the channel. Diffusion in the granular hydrogels was thus fast compared to the nanoporous system. To quantify rates of diffusion within the bulks of the hydrogels, normalized intensity profiles were fitted to Fick’s 2^nd^ law of one-dimensional diffusion (Eqn 1) to approximate diffusivities within the different systems (Supplemental Figure S3). To approximate the diffusivity (*D*∞) of albumin the Stokes-Einstein hydrodynamic radius (Eqn 2) was used:

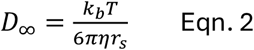

where *k*_*b*_ is Boltzmann constant, T is absolute temperature, η is the solvent viscosity, and *r*_*s*_ is the solute hydrodynamic radius. In water at 20°C the diffusion coefficient of albumin was calculated to be 59.7 μm^2^/s (Figure 6D, grey dashed line).^66^ The diffusion coefficient of FITC-albumin observed in the bulk NorHA gel was 0.02±0.007 μm^2^/s. The diffusion coefficient of FITC-albumin in the NorHA granular hydrogel were observed as 101.3±7.84 μm^2^/s and 187.4±16.8 μm^2^/s for annealed and unannealed granular systems, respectively (Figure 6D). There was a significant difference in the FITC-albumin diffusivities observed in the bulk and both granular hydrogels diffusion coefficient as expected (Figure 6D). There was also a modest, but significant increase in the diffusivity observed within the granular hydrogel when microparticles were left unannealed within the bulk compared to when they were annealed. The modest increase in diffusivities in the granular systems compared to the ideal solution is attributed to slight convective flows across the granular hydrogels that could not be eliminated with during experimentation, which are discussed further below.

**Fig 6.**
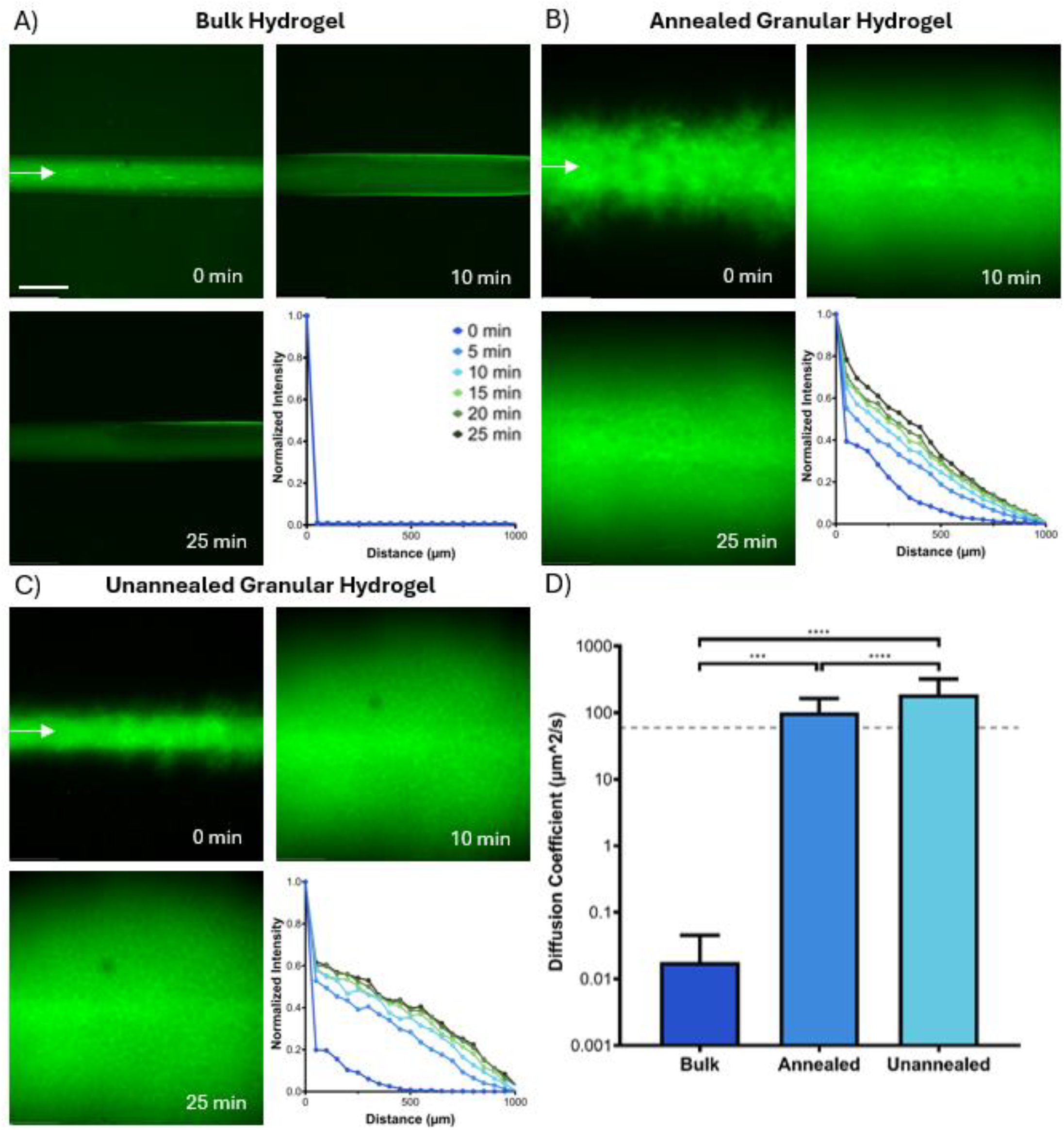
Representative images of FITC-BSA diffusing into the hydrogel from the channel at 0, 10, and 25 mins for A) a bulk nanoporous hydrogel, B) a fully annealed granular hydrogel, and C) a granular hydrogel with the microparticles directly adjacent to the channels are crosslinked, along with the intensity profiles of the fluorescent images that where then used to calculate the diffusion coefficients (scale bar 500µm). D) Calculated diffusion coefficients of bulk, annealed granular, and unannealed granular hydrogel, showing that the nanoporous bulk hydrogel diffusion is slower than the granular hydrogel that is microporous. A statistically significant different between all hydrogels diffusion coefficient was observed. The grey dash line is the diffusion coefficient of BSA in water. (p-values are *** p<0.001, and **** p<0.0001)

In addition to NorHA-based microparticles, PEG-Nb microparticles were used to form granular hydrogels to consider the potential for the device to be used and channels to be formed with multiple hydrogel types and particle sizes. A generalizable approach would allow microenvironmental conditions to be defined using a variety of hydrogel formulations. PEG-Nb microparticles were also formulated using a phase separation process, rather than emulsification, that resulted in particles with a diameter of 12.37±6.76 µm (Supplemental Figure S2). When the device was loaded with PEG-Nb microparticles and selectively crosslinked along the channel, stable channels were observed. As above, this was confirmed by continuous flow of medium through the channels that contained fluorescently labeled solutes.

To observe the potential to establish complex, multi-solute gradients within granular hydrogels, a granular hydrogel with two channels was formed within the device. RhodB, a small fluorescent molecule, was exhibit rapid diffusion through the device, given its small size. RhodB was introduced into the top channel in the device in a flow from left to right and FITC-albumin was introduced into the bottom channel from right to left (**Figure 7**, top row: FITC-albumin in green; middle row: RhodB, pseudo-colored blue; bottom row: overlay of FITC-albumin and RhodB). The differences in diffusivity of the small RhodB molecule and larger FITC-albumin protein were readily apparent. Twenty minutes after the channels were filled RhodB had diffused across the device while the diffusive front of the soluble FITC-albumin was still close to the main channel wall (Figure 7). After 40 minutes both gradients extended from the channel into the middle of the device (Figure 7).

**Fig 7.**
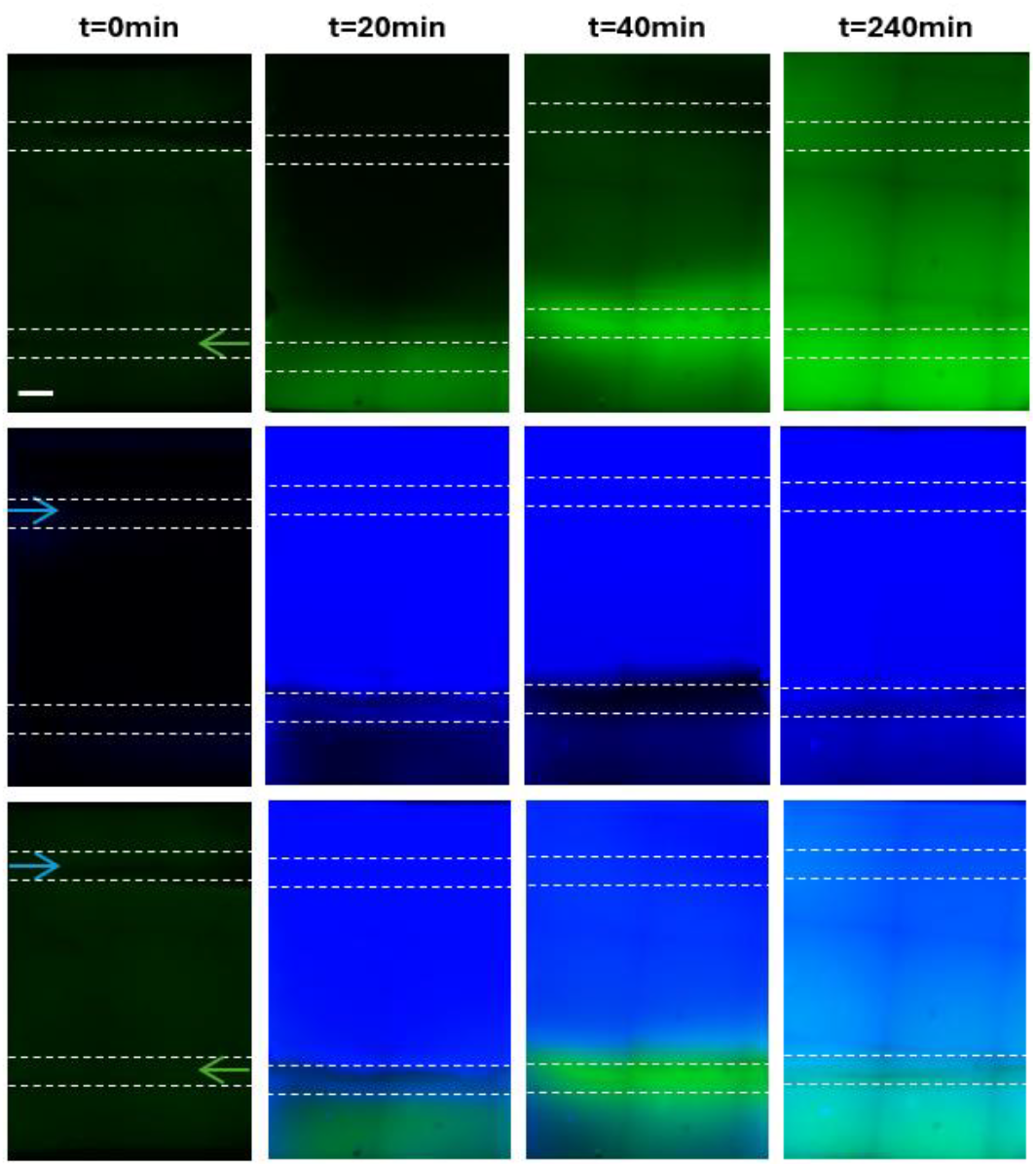
Counterflow diffusion is shown by rhodamine B (blue) and FITC-BSA (green) being flowed through the device over time. Top row is FITC-BSA, second row is rhodamine B, third row is rhodamine B and FITC-BSA overlay, and bottom is rhodamine, FITC-BSA, and brightfield overlay images. (scale bar 1mm).

FMs were perfused through the channel and timelapse imaging allowed for tracking of the FMs as they flowed through the device (Supplemental Videos 1-3). Imaging of FMs revealed flow patterns and differences in porosity between hydrogel formulations. Within the bulk hydrogel FMs were confined to the channel in which they were introduced and were not observed to move into the bulk of the hydrogel. Within the annealed and unannealed granular hydrogel groups, FMs were observed being convected along the channel length and into the surrounding granular hydrogel system. From this experimentation and the diffusion experiments, described above, it was clear that when the medium was introduced into the channel, the change in pressure upon initiating convective flow drove radial flows from the channel in the microporous granular systems. This carried fluorescent solutes, above, and FMs, here, with the radial flow. The pore spaces between microparticles within granular hydrogels were thus able to support convection through the bulk of the material.

### 3D Printing

To establish the ability to use 3D printing processes to define construct composition in a microdevice where it is also possible to control gradients of soluble factors, we printed into unannealed microparticles in the bulk region of the granular hydrogel between channels prior to closing the device. This was possible in the systems where crosslinking only occurred between microparticles adjacent to the channels, allowing the granular hydrogel to flow around a needle in an embedded 3D printing process (**Figure 8**). To demonstrate the potential of 3D printing in defining material heterogeneity, discrete voxels were printed in the center region of the device. According to the computer design and print parameters, the gelatin microparticles voxels that were printed are about 3 mm in diameter. Voxels were positioned with 5 mm between centers (1.5 mm space between each voxel) and 2.5 mm away from the PDMS wall around the device’s center region. The voxels were aligned within the center of the device, 3 mm from the channels on either side. Despite irregularities in voxel circularity, spacing and alignment were well-controlled.

**Fig 8.**
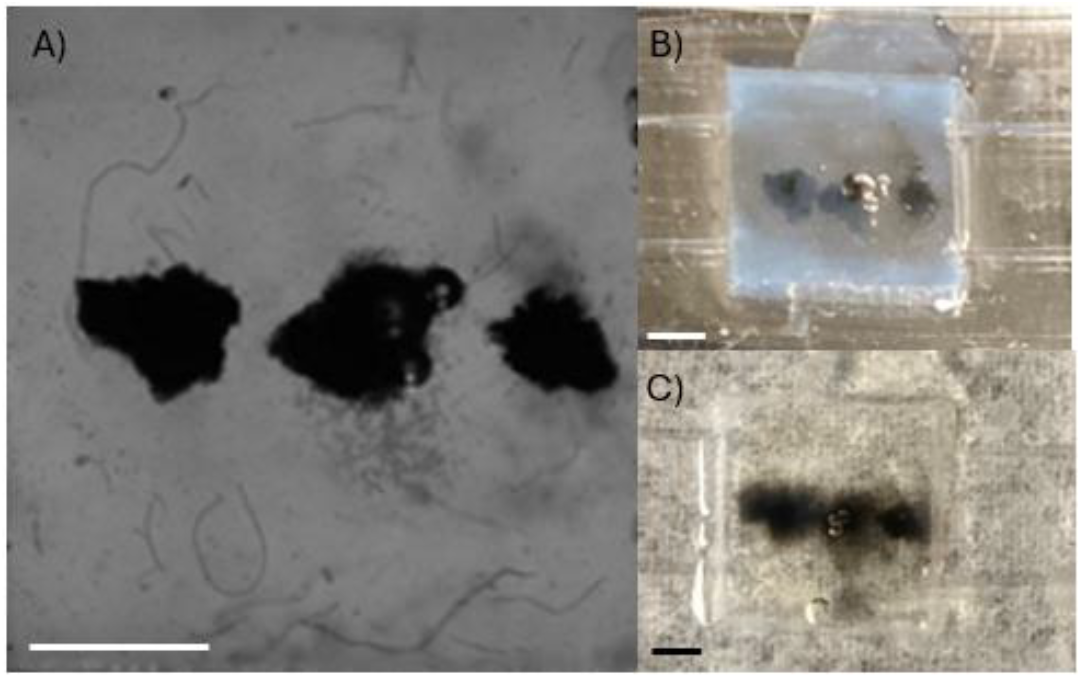
3D discrete printed gelatin microparticle voxels within unannealed microparticles within the device between the channels A) brightfield image of the print in the device, B) the device right after printing, and D) one hour after printing and the gelatin microparticles beginning to melt in the device. (scale bar 1mm).

## Discussion

Advances in microphysiological systems and *in vitro* tissue models will benefit from systems where it is possible to establish complex material compositions of material, and control critical soluble and matrix bound signals. Granular hydrogels facilitate the design of material complexity that is not available in continuous bulk hydrogels through inherent features like microporosity and their discrete particulate nature^39^ as well as compatibility with embedded printing processes^67–69^. The development of a microfluidic system that integrates channels within granular materials will allow control of soluble signals within granular systems, annealed or unannealed, and will allow studies in granular hydrogels without unannealed microparticles dissociating into surrounding media. Additionally, a device compatible with 3D printing opens new possibilities for controlling combined soluble and physical signals.

Granular hydrogels were formed from norbornene-modified hyaluronic acid (HA) and polyethylene-glycol (PEG) microparticles (Figure 2 and Supplemental Figure S2). Stoichiometric control of norbornene functionalization has been shown to allow design of physical and biochemical functionalities in hydrogels ^64,70^ and to facilitate interparticle crosslinking in granular systems.^71,72^ This control was leveraged here to design systems with excess norbornene groups to facilitate interparticle crosslinking adjacent to channels to stabilize granular hydrogels around the channel lumen, while leaving the bulk of the granular system uncrosslinked. Packed microparticle systems that lack interparticle crosslinking retain the ability for individual microparticles to move past one another, preserving inherent stress yielding properties within granular materials that can be challenging to achieve without careful molecular design in continuous bulk hydrogels (Supplemental Figure S4). These dynamic behaviors can be seen in the rheological shear thinning and self-healing responses of the uncrosslinked granular systems (Figure 3A). These properties are desirable in allowing bioprinting to control material architectures^60^ and in permitting dynamic cell behaviors stress yielding behaviors^73^. However, as shown here, unannealed granular systems fully dissociate in culture environments without annealing (Figure 3B) in the absence of a microdevice platform.

To address the dissociation of unannealed microparticles and enable unannealed systems to be maintained in culture environments, we developed a device to contain granular hydrogels in fluidic environments. Here, a PDMS device that contains microchannels within the granular system was developed to allow for introduction of controlled flows. Critically, this work also developed the ability to spatially control the annealing of microparticles to enable microparticles that form the channel wall to be stabilized by interparticle crosslinked while leaving the bulk of the granular system unannealed. The approach here demonstrated that localizing the photoinitiator to a small volume of solution that wet the channel forming needles results in stable crosslinking of channels around the needles. A similar diffusion-based strategies has also been used to spatially controlled crosslinking during 3D printing.^74^

Stable channels withing anneal and unannealed granular systems were observed to support flow and remain open throughout the duration of the experiments that leveraged the channel structures to perfuse constructs and deliver soluble compounds. Channels were successfully templated around a needle (Figure 5, grey dash line), with diameters within hydrogels becoming smaller due to swelling of the hydrogels after needle removal. Swelling was observed to increase as total crosslinking within the hydrogels decreased, as would be expected. Data showed a trend where continuous hydrogels exhibited the largest diameter channels and unannealed granular hydrogels the smallest. The granular hydrogel systems used are also expected to be compatible with processes for creating channels using 3D printed inks designed to be printed into granular systems and removed to leave vessel like structures after inducing annealing.^52,75–77^

Controlled gradients within granular hydrogels were demonstrated by delivering molecules that diffused from the channels into the granular material. Diffusion of FITC-albumin within the granular hydrogels established a gradient of a soluble protein between two channels where one channel served as a source of the protein and the other as a sink. Notably, this gradient was established quickly, with diffusive transport occurring with diffusivity similar to that seen in free solution. This should allow for design of spatially and temporally controlled signaling gradients, which are important in MPSs. In quantifying the diffusion coefficient of FITC-albumin an increased diffusivity was observed compared to the diffusivity in water (Figure 6 D, iv, gray dashed line).^66^ In studies here, this was attributed to convection of the FITC-albumin solution from the channel into the surrounding hydrogel. The interstitial space between microparticles is likely to provide little flow resistance, and this slight imbalances in pressure as flows were initiated resulted in convection that led to a calculation of apparent diffusivity values exceeding the expected value in free aqueous solution. Moving beyond this first generation of microfluidic-granular hydrogel system, refinements to control for introducing flows, to controlling pressure gradients, and device packaging will be pursued.

In physiological systems, complex extracellular signals are integrated through cell signaling pathways to determine cellular behavior. To assess the potential to define complexity in soluble signals, here the diffusion of multiple soluble compounds across a granular system established opposing concentration gradients of two model molecules, RhodB and FITC-albumin (Figure7A). Each channel within the device acted as the source for one compound and the sink for the other. Additionally, towards demonstrating the potential for using diverse granular hydrogel materials with a device, we considered small (∼10 µm in diameter) PEG-Nb microparticles in addition to larger NorHA particles. Our results suggest channels within granular hydrogels could be used for a range of microparticle sizes and for both naturally derived and synthetic polymers.

It is expected, based on the known microporosity within a granular hydrogel and the calculations of apparent diffusivity, that the granular hydrogels support interstitial flows. Among disease models and physiological systems, interstitial flows are important in processes including cancer metastasis^7^ and microvascular network formation^18,21,28^. The presence of radial flow into the granular systems’ interstitial space was directly observed through the inclusion of FMs (Supplemental Videos 1-3); these small particles were carried in suspension in flow. When the pressure differences between channels resulted in flow across the granular hydrogel, FMs were seen to flow into the material’s interstitial space. Because of their large micron-scale size relative to molecular solutes, convection of FMs suggests the potential for cellular movement within such a granular hydrogel, highlighting the biological attraction of these materials and the need for culture systems that allow researchers to exploit these features in designing synthetic cellular matrices.

Finally, this system allowed 3D printing to be used to design material heterogeneities within the granular hydrogel on the microdevice. Controlling crosslinking of microparticles to one another adjacent to the channels allowed for the granular hydrogel between the channels to be used as a support bath for 3D printing. A demonstration of control over material composition was shown here in Figure 8. The simple, but specific and regular computer-guided placement of the depots of a granular gelatin material could be extended to include materials with different mechanics, bio-functionalities, and porosities. Additionally, bioprinting will help create biological systems that have controlled placement of cells and materials within the microfluidic device, which will be valuable for reproducible control of biological environments and emergent structures. The system thus presents opportunities to control a range of cell-instructive cues not available in other systems. Consequently, it is expected that a device that allows application of controlled flows to granular systems can be support well-defined biomaterial systems *in vitro* and the development of more sophisticated models for tissue engineering and microphysiological applications.

## Conclusions

In this work, a microfluidic device was developed to contain granular hydrogels, with the ability to selectively crosslink microparticles adjacent to channels within the device, leaving microparticles in the hydrogel volume between the channels uncrosslinked to one another. These unannealed granular hydrogels have valuable properties but are otherwise challenging to be used as scaffolds for tissue engineering or microphysiological systems without undergoing erosion into culture media. Here, a device that facilitated easy loading of packed microparticles allowed the fabrication of channels into which media could be introduced, without scaffold erosion. Through spatial control over crosslinking that locally stabilized channels, gradients of soluble compounds could be established across the hydrogels. Additionally, the uncrosslinked hydrogel could support material to deposition via 3D printing. The combined microfluidic device and fabrication techniques demonstrated here allow for spatiotemporal control of soluble and material-based signals in dynamic granular systems. The system would allow increased complexity to be designed into biological systems *in vitro* for a variety of biomedical applications.

## Data availability

The data supporting these findings is available in ESI. Additional data is available upon request from the corresponding author.

## Acknowledgements

This research was supported by supported by NIGMS R35 GM147410.

