## Supplemental figures for "Integrating Microchannels and Flows into 3D Printable Granular Hydrogel Matrices"

### Supplemental Material

#### Rheology Methods

Bulk gel *in situ* rheology was performed, the rheometer was equipped with a 20 mm sandblasted geometry and UV light at 100 mW/cm^2 was used. For the gelation curve a time sweep (1 Hz, 1% strain) was performed. Frequency sweeps (0.1-100 rads/s, 1% strain) were used to assess the storage and loss moduli of the hydrogel.

#### Example G-code

```
G90 ; use absolute coordinates
M83 ; use absolute distances for extrusion
G1 Z5; lower 5mm
G1 E1 F10; extrude 1mm at 10mm/min
G4 S2 ; pause 2 seconds
G1 Z5 F50 ; raise the needle
```

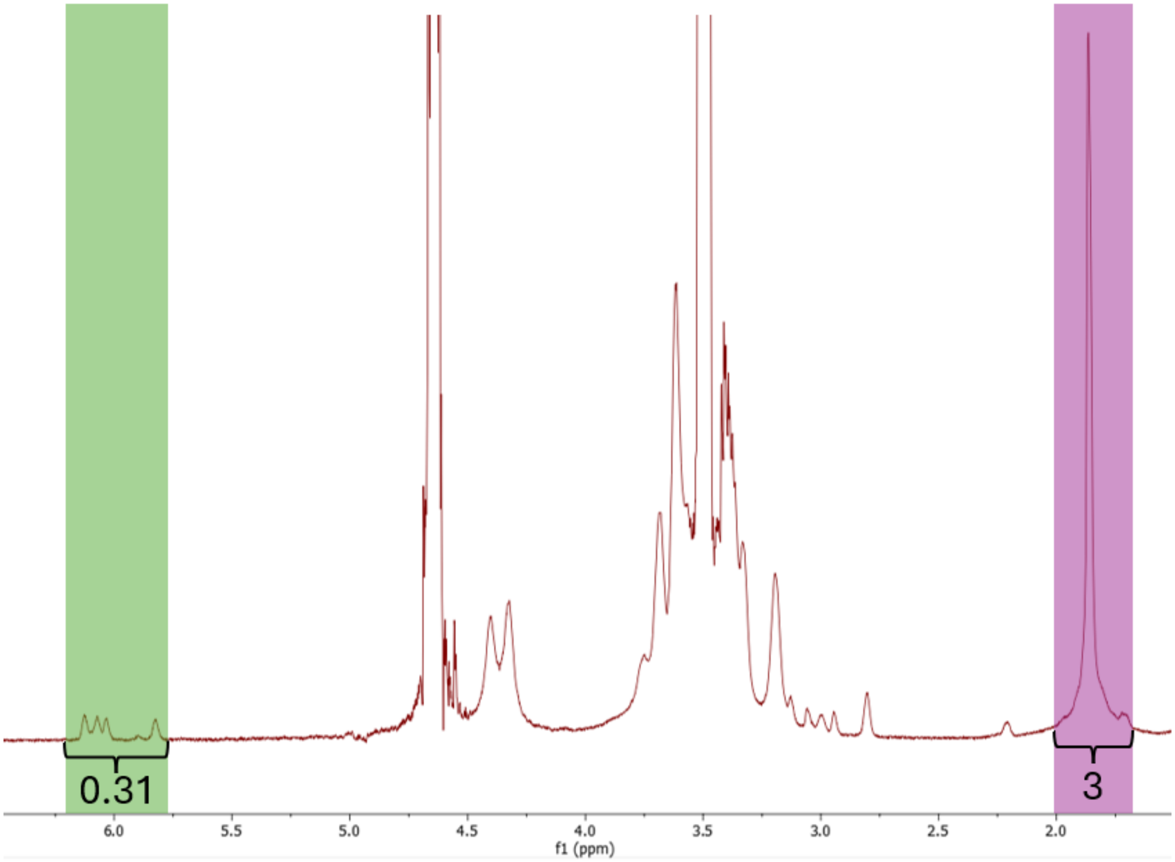


**Figure S1** Norbornene-functionalized hyaluronic acid (NorHA) 1H NMR spectrum. The resultant NMR spectrum above illustrates a modification of 15.5%.

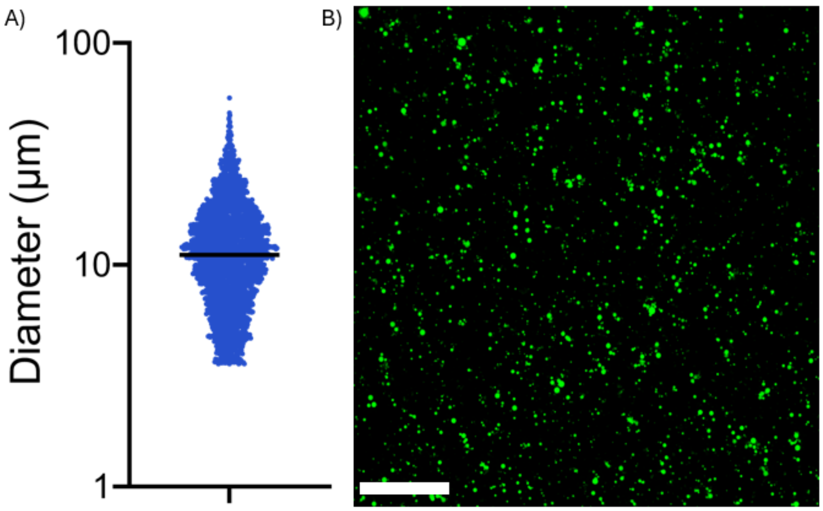


**Figure S2** A) Size distribution and B) representative image of PEG-Nb hydrogel microparticles (scale bar 500μm).

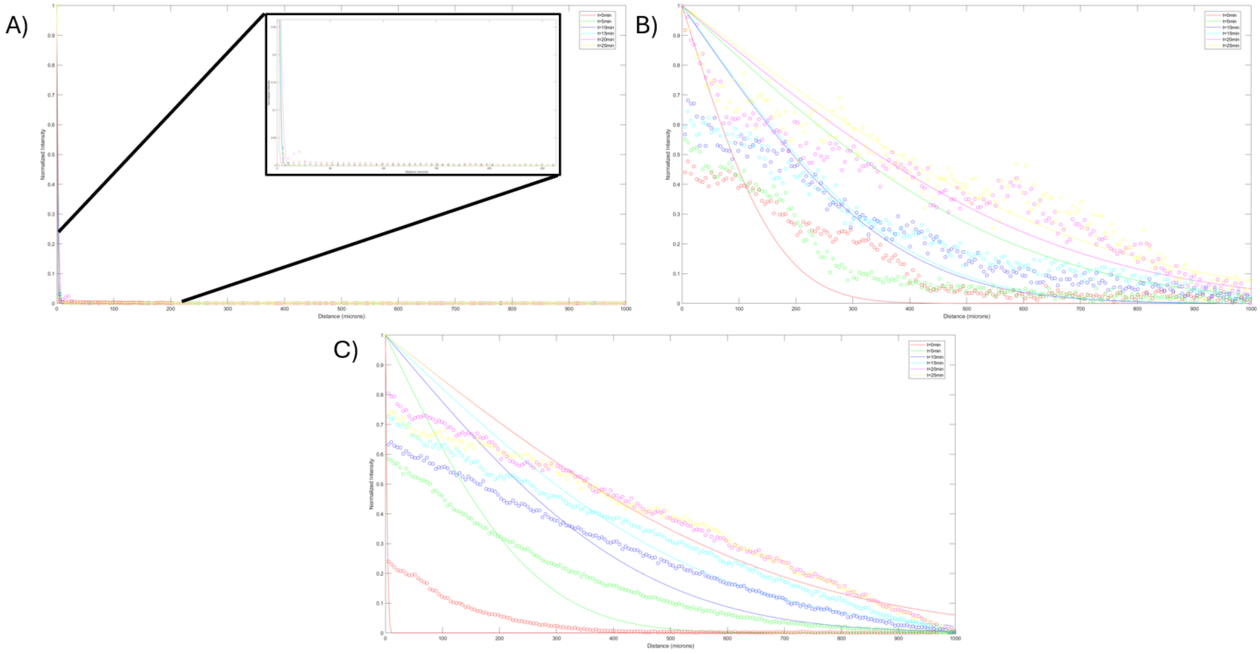


**Figure S3** Intensity profiles used to quantify the diffusion coefficient of FITC-BSA, raw data (circle) and fitted curve (line) of A) bulk (zoomed in graph of bulk gel fit (insert)), B) annealed, and C) unannealed hydrogels.

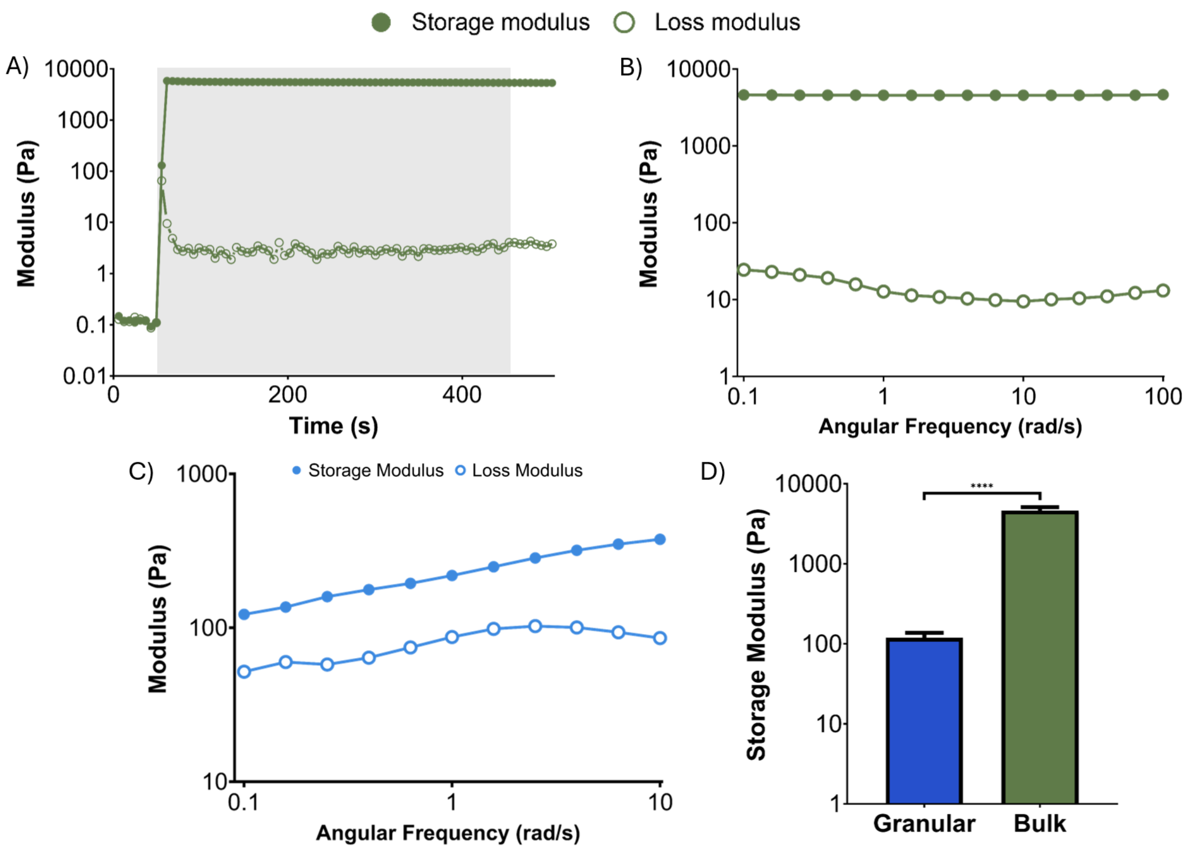


**Figure S4** A) NorHA gelation curve, grey area represents the light on. B) Bulk NorHA gel frequency sweep. C) NorHA granular hydrogel frequency sweep. D) Storage Modulus of NorHA granular hydrogel and bulk NorHA gel. (**** $p<0.0001$)
